# Pandemic Legacy: Medical Facemasks as a Potential Source of Marine Microplastic?

**DOI:** 10.1101/2025.10.13.682204

**Authors:** Olivia Dillon, Ina Benner, Tatiana Zaliznyak, Carolina Cisternas-Novoa, Janika Reineccius, Gordon T. Taylor, Uta Passow

## Abstract

Understanding the main sources of microplastic pollution is key towards developing efficient measures to reduce microplastic loadings to marine waters. Yet identifying the main sources of marine microplastic is challenging. Source tracking should be easier in marine bays where inputs are limited. In 2021 we determined the concentrations and characteristics of microplastics > 300 μm in surface waters of Placentia Bay, Newfoundland; a bay with negligible river input in an area of low population density, and no plastic processing plants in the vicinity. Microplastics contributed 2-14% to particulate organic carbon (> 300 μm), and concentrations ranged from 0.11 to 0.67 particles m^-3^, a relatively high level given the region’s low population density. Microplastic diversity was low; fiber and fragment concentrations dwarfed those of other shapes, and polypropylene (PP) dominated, with transparent PP fibers specifically contributing near 50% to the total microplastic inventory. The overwhelming dominance of transparent PP fibers, as well as the exceptionally high proportion of long fibers, suggest that a distinctive input of large, transparent PP fibers overlaid “background” inputs from other sources. A ballpark estimate indicates that weathering of medical facemasks used during the COVID-19 pandemic are a likely explanation for the dominance of transparent PP fibers in Placentia Bay in 2021. Similar inputs may have affected many other aquatic environments globally, but might not have been observable in systems where other continuous input pathways are high.

## Introduction

Microplastics, which include plastic particles smaller than five millimeters in size (1) are characterized by their polymer composition, shape, size, and sometimes color. They are of growing concern, mainly because of their pervasive presence, effects, and persistence (2–4). A prerequisite for the development of effective strategies to reduce marine microplastic pollution, is the identification of the main sources and transport pathways (5–7). Only once we understand the main pollution pathways of microplastic, can effective alterations to plastic production and waste management be implemented.

Generally, most plastic in the ocean (80%) originates from land-based sources such as landfills, sewage, cities, including cars, and agriculture (8, 9). Correlations between microplastic abundances and coastal human population centers, rivers or coastal industrial sites are well established (10). Fishing and aquaculture activities also release a large amount of microplastic (11, 12) that contribute about 18% to marine plastic debris worldwide (9). Correlations between concentrations of polyethylene (PE) and PP fibers and fishing activity have been observed resulting from fishing nets and lines shedding synthetic fibers during use (13) and continuous release of microplastics from improper disposal and accidental loss of fishing equipment (14) (15–17). However, with the exception of certain niche-use polymers, whose limited applications reveal their origin, it is usually challenging to assign microplastic particles in the marine environment to specific sources (18).

Nevertheless, geographical differences in microplastic types (shape, polymer composition) (19) point to the importance of local sources and waste management strategies, as well as to the hydrographic conditions in determining the microplastic composition in bays and fjords (12). This suggests that some progress can be made in such systems by looking at the composition of inventories in relation to possible local source sites and currents. For example, in a comprehensive study of microplastic particles > 300 μm in the Mediterranean’s surface waters (20), the authors attributed regional differences in the relative composition of microplastic between basins to differences in input from rivers and increased shipping activity.

Identifying specific sources is potentially more successful in bays with a limited number of input sources, e.g., far from cities, traffic, and with minimal river contribution. The present study, which was part of the Coastal Environmental Baseline Program initiated by the Department of Fisheries and Oceans (DFO), Canada, aimed to determine potential sources and distribution pathways of marine microplastic in Placentia Bay. Placentia Bay, located along the southeast coast of Newfoundland and Labrador, characterized by insignificant river-run-off, the absence of coastal communities exceeding 5000 people, and the absence of plastic processing plants. Thus, a comparatively minimal input of microplastic particles from land-based sources was expected. As Newfoundland is relatively remote, aerial input is assumed negligible. Marine activities were hypothesized to contribute moderately, because Placentia Bay has supported commercial and recreational fishing industries for decades (Government of Canada, 2018).

While, we initially anticipated microplastic concentrations in Placentia Bay to be lower than in most surveyed bays, and expected source tracking to be feasible due to limited expected inputs, sample collection in the summer 2021 occurred shortly after the exponential rise in the release of medical face masks into the environment (21). Estimates suggest that in Canada alone > 24 million facemasks were discarded daily (22), many of which were disposed improperly, in part because of the fear of contamination (21). Common types of disposable face masks mainly disintegrate into PP particles (21), specifically, transparent PP fibers (23–26). We thus hypothesized that PP fibers (stemming from face masks) would be over-represented in Placentia

Bay surface waters in 2021. The assumed low background input of microplastic into Placentia Bay would make such a potential input of PP fibers during 2020 observable.

## 1. Materials and Methods

### 2.1 Study Area and Sample Collection

Microplastic samples were collected from the R/V Port Tender Royal Breeze at nine stations in Placentia Bay, Newfoundland (47.25 N, 54.25 W; **Figure 1**). At the mouth, the outer Bay is wide and relatively deep (80 km, > 250 m), with a deep channel stretching along the eastern side of the Bay, which is separated from the western side by several islands. The enclosed index of 0.5-0.6, calculated from the product of the square root of the bay’s surface area (a) and its maximum depth (d_max_) divided by the product of width (w_m_) and depth (d_m_) at the bay’s mouth (27), suggests a short residence time of water in Placentia Bay. However, the islands located in the centre of the bay (**Figure 1**) protect the waters in the inner bay (28), leading to a higher water residence times there.

**Figure Caption 1:**
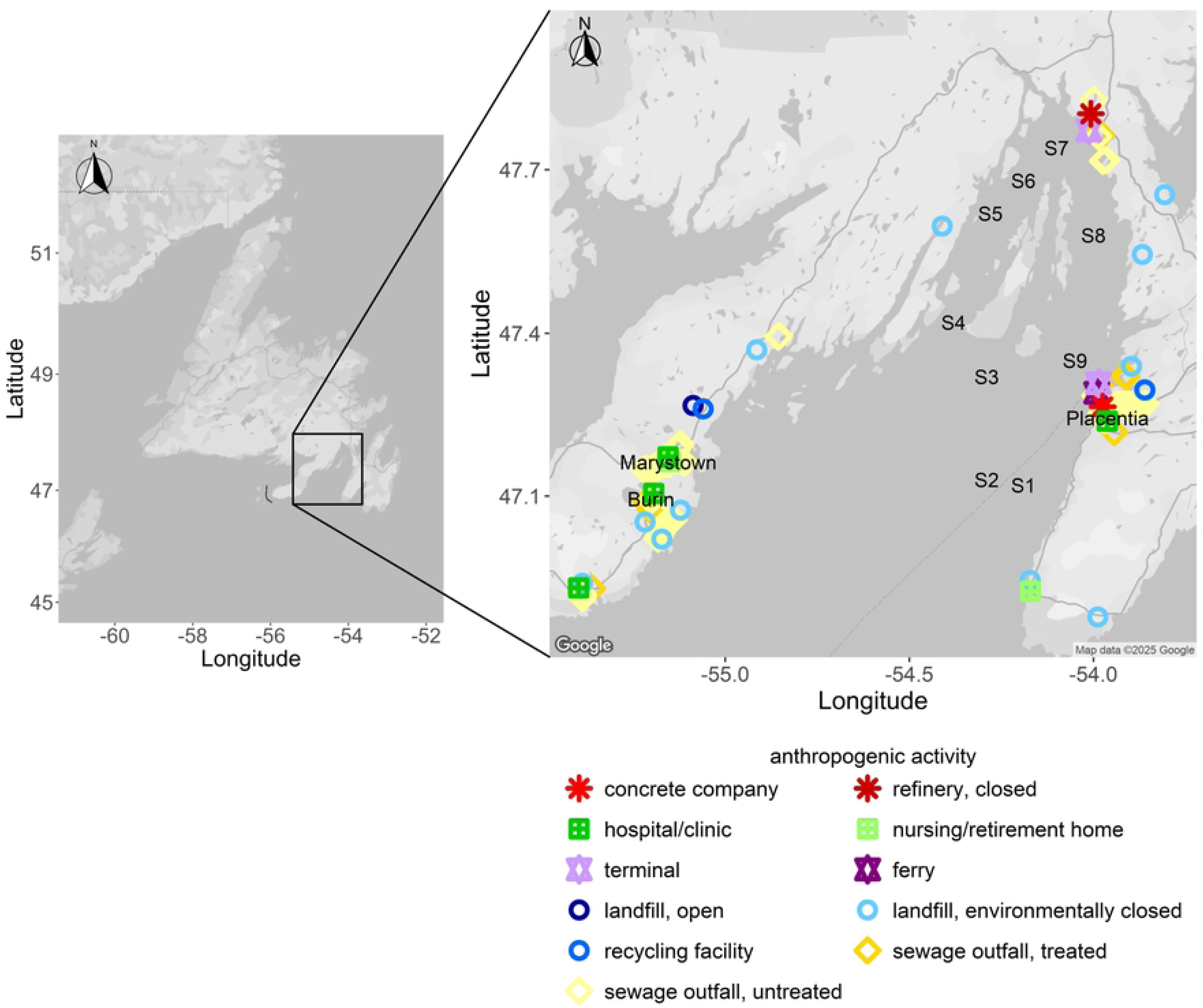
Map showing Placentia Bay on the southern coast of Newfoundland, with sampling locations S1-S9 marked. Also depicted are sites of potential interest as terrestrial microplastic sources.

All samples were collected during the summer of 2021; stations S1 to S4 in July and stations S5 to S9 in September. Station locations chosen to represent the whole Bay, were categorized according to their position as outer Bay stations S1 and S2, central Bay stations S3, S4, and S9, and inner Bay stations S5, S6, S7, and S8 (**Table S1**). Surface water temperatures ranged between 15 and 18 °C, and surface salinity varied between 34 and 35 PSU, measured with an EXO2 multiparameter sonde (YSI Inc., USA).

The coastal zone around the Bay includes three towns, Placentia, Marystown, and Burin, each with populations between 2000 and 5000 people, as well as some smaller communities.

Potential microplastic source sites (**Figure 1**) include landfills, sewage outfall sites, an industrial harbor/ferry terminal at Argentia, health care facilities, an oil refinery, and a concrete manufacturing company, with one cluster of sites on each coast, and another at the top of the Bay, associated with the oil refinery (**Figure 1**).

At each station, samples were collected with a neuston net (300 µm mesh and a 50 x 100 cm opening) at the sea surface and additionally a plankton net (300 µm mesh, 75 cm diameter opening) at ∼2-5 m depth, allowing a comparison between both net types (**Supplements, Fig.**

**S1)**. Since the collected microplastic were not statistically different between nets, we treat them as replicates and present the data as averages. The volume of water filtered was measured with a flowmeter (General Oceanics Inc, model 2030R), and nets were towed for one hour at 2.5 knots. Filtered volumes per tow ranged between 1420 and 2250 m^3^ for the neuston net and between 1120 m^3^ and 1480 m^3^ for the plankton net. Volume calculations for the neuston net assumed that 20 % of the net mouth was above the water line, a modest assumption, resulting in concentration estimates at the low end of the range. Sampling with two nets per station reduced sample collection bias and addressed patchiness. Samples were transferred into 2 L Mason glass jars and stored at 4 °C. A 6 mL sub-sample was removed for immediate particulate organic carbon (POC) analysis, and the remaining sample was stored for two to three months until microplastic analysis.

### 2.2 Particulate organic carbon analysis

Subsamples (6 mL) from stations S5 to S9 were split into three 2-mL replicates, and particles were filtered on pre-combusted (400 °C, 6 h) glass fiber filters (25mm, Whatman GF/F). Dried filters (60 °C, 24 h) were exposed to acidic fumes for 24 hours to remove inorganic carbon, and then dried again (60°C, 24 h). Afterwards, filters were pelleted and analysed with a CHN analyzer (PerkinElmer, 2400 Series II CHNS/O).

### 2.3 Sample preparation for microplastic analyses

To prevent contamination, only glass or metal implements were used during sample collection, storage, processing, and analysis, with the exception of the nets (mostly nylon). Natural fiber clothes, including cotton masks, were worn during field and lab work, and fibers of synthetic foul weather gear were collected as controls, using sticky tape. Additionally, all laboratory procedures were conducted in a plastic-free lab under a designated fume hood. Blanks were generated daily for each process and sample by placing a glass petri dish filled with Ultrapure Milli-Q water in the work area. All particles in blanks were analyzed with Raman microspectroscopy.

Each net tow sample was sieved onto a 300 µm stainless steel mesh, rinsed with Ultrapure water, and the retentate was freeze-dried for 3-5 days to remove all moisture (29). Biogenic organic material was digested for 24 hours at 50 °C with 50-230 mL of hydrogen peroxide (H_2_O_2_ 30%, Sigma), depending on sample volume. Then an equivalent volume of acetic acid (24%, Sigma) was added, digesting for another 12 hours at room temperature (29). A gravimetric recovery test using microplastic derived from weathered plastic collected at a local beach confirmed that the digestion method did not affect microplastic recovery. After digestion, the samples were rinsed again onto a 300 µm stainless steel sieve to remove partly digested organic matter and reagents (30). Particles retained on the sieve were rinsed off with DI water and captured on polycarbonate membranes (10 µm pore size, 47 mm diameter), using several filters where necessary. The sieve was examined microscopically to ensure no microplastic remained.

Macroscopically visible particles were isolated manually into glass scintillation vials using forceps and enumerated as described below, as their presence on the filter made identification of smaller particles difficult. If filters retained too much organic detritus, which interfered with reliable enumeration of the microplastic particles, a second digestion step was conducted by adding a small amount of 30% H_2_O_2_ and, after 24 hrs adding the same volume of 24% acetic acid, before filtering particles onto a new filter. Both, the new and original filters were enumerated. All filters were stored in glass petri dishes for later enumeration and identification.

### 2.4 Microscopically determined microplastic abundance

Particles visually identified as microplastic were counted and categorized by shape, size, and color under a stereomicroscope (Wild Heerbrugg 256530, at 75x) on all filters. Particles were identified as plastic based on color (e.g. bright colors) and rigidity, assessed by probing with forceps. Particles that were soft or broke apart upon probing were excluded.

Four morphotypes of microplastic were distinguished; fibers, fragments, films, and microbeads (31). Fibers were identified by a length-to-width ratio of > 3 and were sized by length (mm) (32). Tangled fibers, which made up roughly 10-20% of fibers were separated using dissecting tweezers and then individually counted and measured.

We identified fragments by a length-to-width ratio ≤ 3, with irregular shapes, characterized by jagged edges and uneven surfaces, which distinguishes them from other microplastic forms (33). Fragments were sized by area (mm^2^), as this is faster and more accurate for odd-shaped particles, and the area-based sizes were converted to an equivalent square length. We found only minimal amounts of both films and beads in our samples. Films are defined as a thin layer or sheets of synthetic polymers, commonly found in shopping bags, packaging materials, and agricultural films (34). Beads are spherical plastic particles. Each particle type was categorized into one out of five size classes, using logarithmic scale binning. Increasing bin width with increasing size improves the probability of capturing larger, rarer particles and strengthens counting statistics. All microplastic particles were also categorized into eight color categories: transparent, black, green, white, blue, red, yellow, purple, and orange. On average (± 1 SD), 1190 ± 840 particles were counted at each station and net tow, implying a statistical counting error of 6% (35). As concentrations based on plankton and neuston net samples were not statistically different (**Supplements, Figure S1**), concentrations from both nets are presented as averages.

These visual counts were corrected based on Raman results to account for overestimates during visual identification (**Supplements, Figure S2).** These corrections (see below) offset potential misidentification during visual surveys (36). Thus final, Raman-corrected concentrations are considered to be low estimates, because microplastic particles are easily lost during processing, as many polymers adhere to glass surfaces and are not washed off even with careful rinsing. We were not able to correct for those losses.

Particle size distributions are presented as differential number size distributions dN/dl (# particle m^-3^ mm^-1^), where N equals the number of particles in a given size class, and l the corresponding length interval. The best fit power law relationship dN/dl = *A* l^-*b*^, where *A* and *b* are constants, was calculated separately for fibers and fragments. A theoretical *b* value of 4 suggests equal particle volumes in each size class, while smaller *b* values indicate a higher proportion of large particles (37, 38). As is common practice, our *b* value was calculated omitting the smallest size class, where concentrations are often artificially low, due to the potential of particles escaping nets or screens (37, 38).

### 2.5 Raman microspectroscopy and corrections of microplastic concentrations

Selected particles were investigated with a Renishaw® inVia™ confocal Raman microspectrophotometer configured with a modified upright Leica® DM2700™ fluorescence microscope (NAno-Raman Molecular Imaging Laboratory (NARMIL) at the School of Marine and Atmospheric Sciences, Stony Brook University). A He-Ne laser (633 nm wavelength) or diode laser (785 nm wavelength) was used as the excitation light source. Spectra were acquired using a 1200 line/mm diffractive grating which provided spectral resolution of 1.1 cm^-1^.

Laser power and exposure time were individually adjusted for each particle to optimize results. Laser power ranged from 1.01 ± 0.08 to 9.66 ± 2.8 mW (10 and 100% nominal laser power) at the sample using a 50x objective for the He-Ne 633 nm laser and from 0.55 ± 0.05 to 17.85 ± 3.12 mW (10 and 100% nominal laser power) at the sample with the 785 nm diode laser.

Spectra were minimally processed using Renishaw’s® Wire 5.1™ software by polynomial baseline correction. Chemical composition of each particle was determined from its Raman spectrum which was compared to the NARMIL internal spectral reference library of well-characterized polymers. Positions and relative intensities of three to six peaks were sufficient for positive identification of any specific polymer.

Raman microspectroscopy was performed on a total of 195 particles (1.8% of all counted particles). Analysis of 166 of these Raman spectra clearly identified 86 particles as microplastic polymers. Another 33 spectra with easily recognizable peaks were identified based on a more informal visual assessment, which proved correct in 81% of test cases where both a visual and a detailed spectroscopic analysis exist (37 cases). Raman analysis focused on the most common particle types and colors observed visually. However, we also targeted moderately abundant particles and those with variable spectra, such as black fragments, to capture reliable data across different particle types (**Figure S2**).

Raman microspectroscopy confirmed that all particles collected as blanks were cotton or cellulose, implying microplastic contamination was minimal during sample processing. Thus, no blank correction was applied. Raman microspectroscopy further revealed that particles identified as plastic during visual counts fell into three groups; unambiguously plastic, unambiguously non-plastic, and a group of ambiguous particles: About 75% of the visually recognized microplastic particles were positively identified as specific plastic polymers, 6% were positively identified as non-plastic, such as cotton, organic detritus or sulfur and 18% were considered ambiguous, because of spectral interferences from strong dyes. We used these values to correct our microscopic counts to avoid overestimations that commonly occur during visual identification (**Figure S2**). We developed a protocol to estimate *probable* and *conservative* concentrations, with *probable* concentrations calculated from microscopic counts reduced by 6%, e.g., the fraction of non-plastic as identified by Raman. Another 18% (the ambiguous particles) were removed for the *conservative* estimates of concentrations. Results and figures depict *probable concentrations* unless noted.

### 2.5 Microplastic carbon content

Microplastic carbon content was estimated for stations S5-S9 based on microplastic concentration in each size class, the calculated average microplastic volume of particles in that size class, the density of microplastic polymers, and their specific carbon content. For fibers, a cylinder was calculated from the average lengths of each size class, assuming a diameter of 15 μm (15). For fragments, the equivalent square length was converted to an equivalent spherical diameter (ESD), and then the volume of the sphere was calculated. The density of 0.92 g cm^-3^ was assumed to calculate microplastic mass for both PP and PE. The carbon content for PE and PP may be calculated from their molecular formula as 86%, and has been measured as 82% (39), which we used to calculate the fraction of POC > 300 µm that resides in microplastic rather than organisms or detritus.

## 2. Results

### 3.1 Microplastic concentration and characteristics

The average *probable concentration* of microplastic particles > 300 μm ranged between 0.11 and 0.67 particles m^-3^ with a median concentration of 0.28 particles m^-3^ (average: 0.31 ± 0.19 particles m^-3^) (**Figure 2**). Using the upscaling correction introduced by (40), microplastic concentrations with an ESD of 10 - 5000 μm would be 1.6 particles m^-3^ in Placentia Bay.

**Figure Caption 2:**
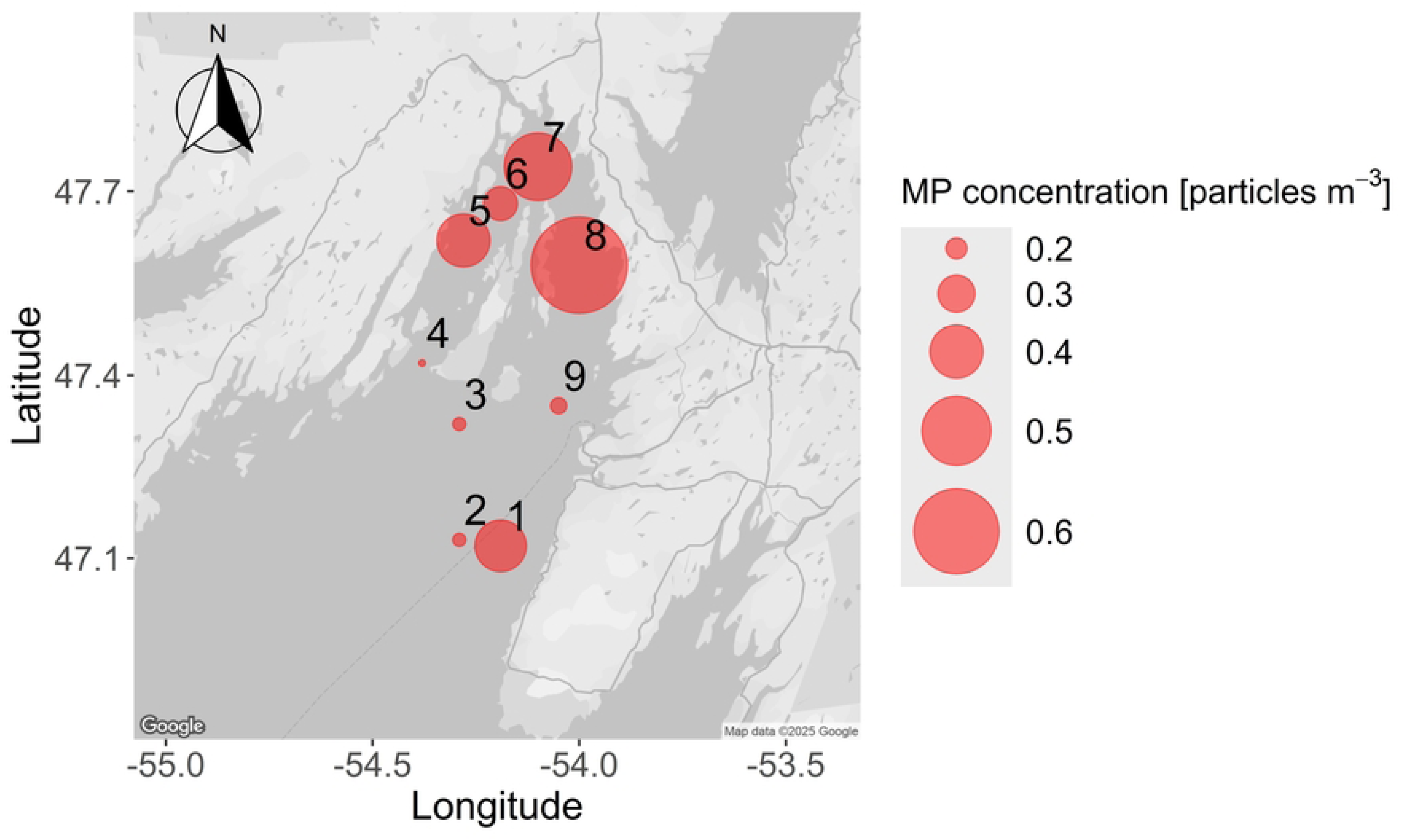
Spatial distribution of probable microplastic concentration in Placentia Bay.

The station with the highest average probable microplastic concentration detected was at the eastern inner Bay station S8 (0.67 ± 0.19 particles m^-3^). The average probable microplastic concentrations at all stations in the inner Bay (S5, S6, S7, S8) were intermediate to high compared to those in the central and outer Bay, likely reflecting increased water residence time in the inner Bay. Average microplastic concentrations at the outer Bay station S2 and all three central Bay stations (S3, S4, and S9) were < 0.2 particles m^-3^. The only exception was the outer Bay station S1 (**Figure 2**), where microplastic concentrations were intermediate (0.4 particles m^−3^), and showed high variability between net tows (**Figure S1**).

Microplastic diversity, based on polymer type, shape, and color was limited to two shapes, fibers and fragments, and four polymers. Fibers dominated, contributing over 80% to microplastic inventories and fragments provided the remainder. Few or no microbeads or films were observed in any of the samples. Microplastic particles consisted of polyethylene (PE), polypropylene (PP), polystyrene (PS), and polyester (PES). Fibers consisted mostly of PP (> 70%) and fragments consisted mostly of PE (> 80%) (**Figure 3**). PE fibers and PP fragments contributed between 10 and 20% each to microplastic inventories. Two fibers and one fragment, all transparent, were PS, and four fibers of different colors were identified as PES. Except for purple, all colors (8) were observed, but transparent particles overwhelmingly dominated (79%) (**Figure 4**). Based on polymer types, shape, and color, we found 26 distinct types of microplastic. Of these 26 types, transparent PP fibers were the single most common type of microplastic, accounting for 47% of the entire microplastic inventory. Transparent PE fragments contributed another 11%, followed by blue PP fibers (6%) and white PE fragments (3%). The remaining microplastic types observed each contributed < 3%.

**Figure Caption 3:**
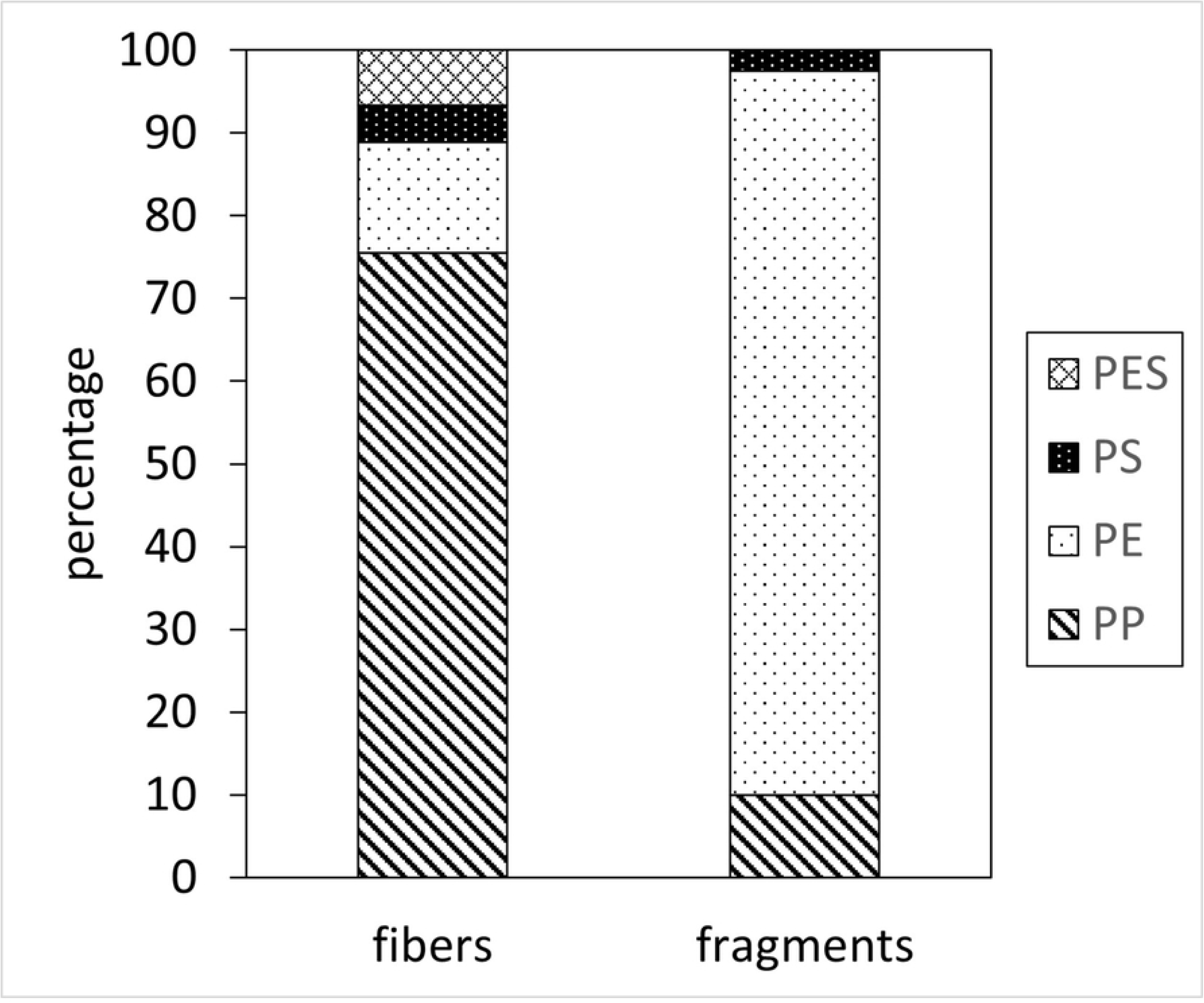
Polymer composition of fibers and fragments analyzed by Raman spectroscopy. Note that only four polymer types were found. Overall, 62% of all microplastic were polypropylene (PP), 26% were polyethylene (PE), 7% Polyester (PES) and 4% consisted of polysterene (PS).

**Figure Caption 4:**
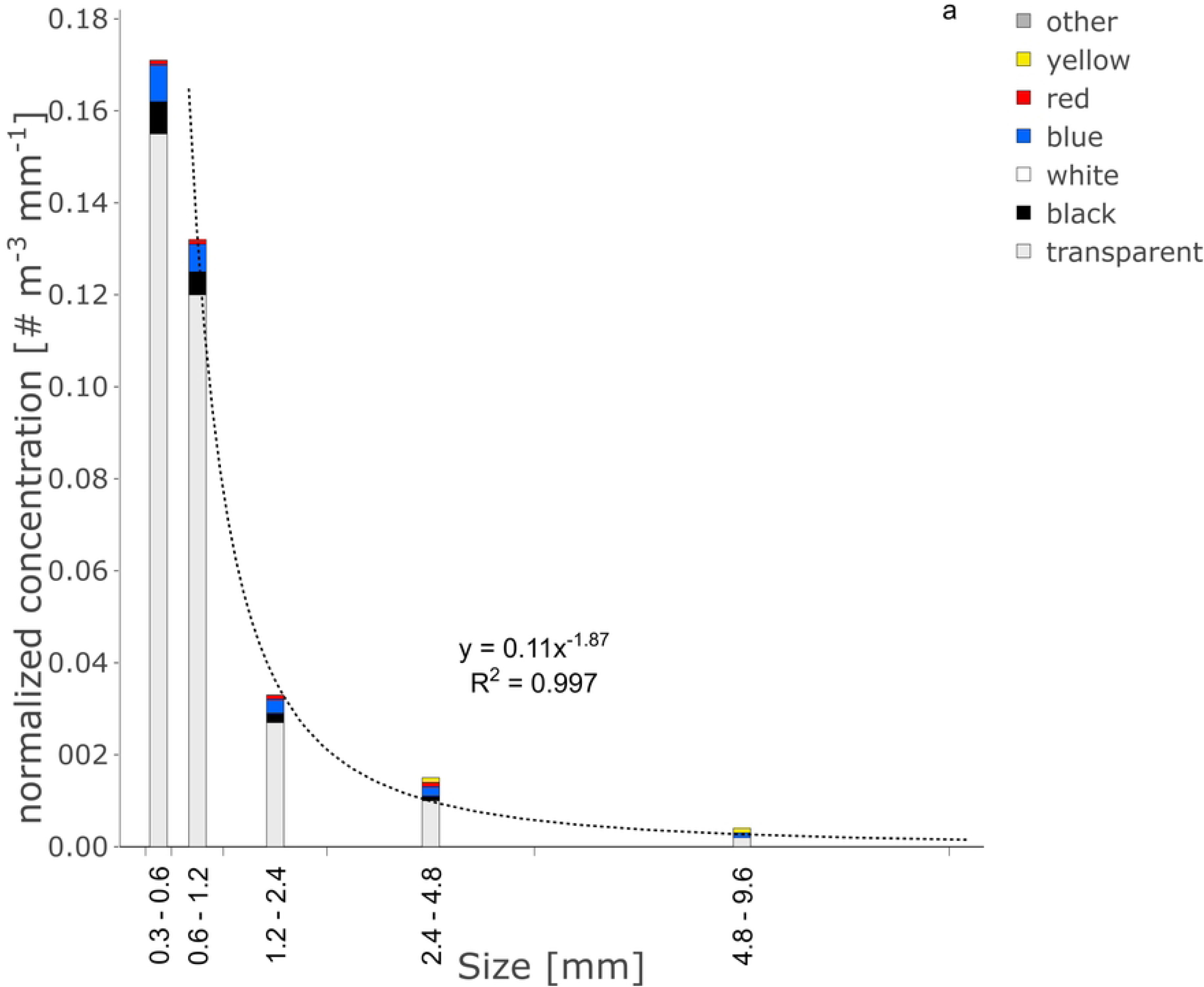
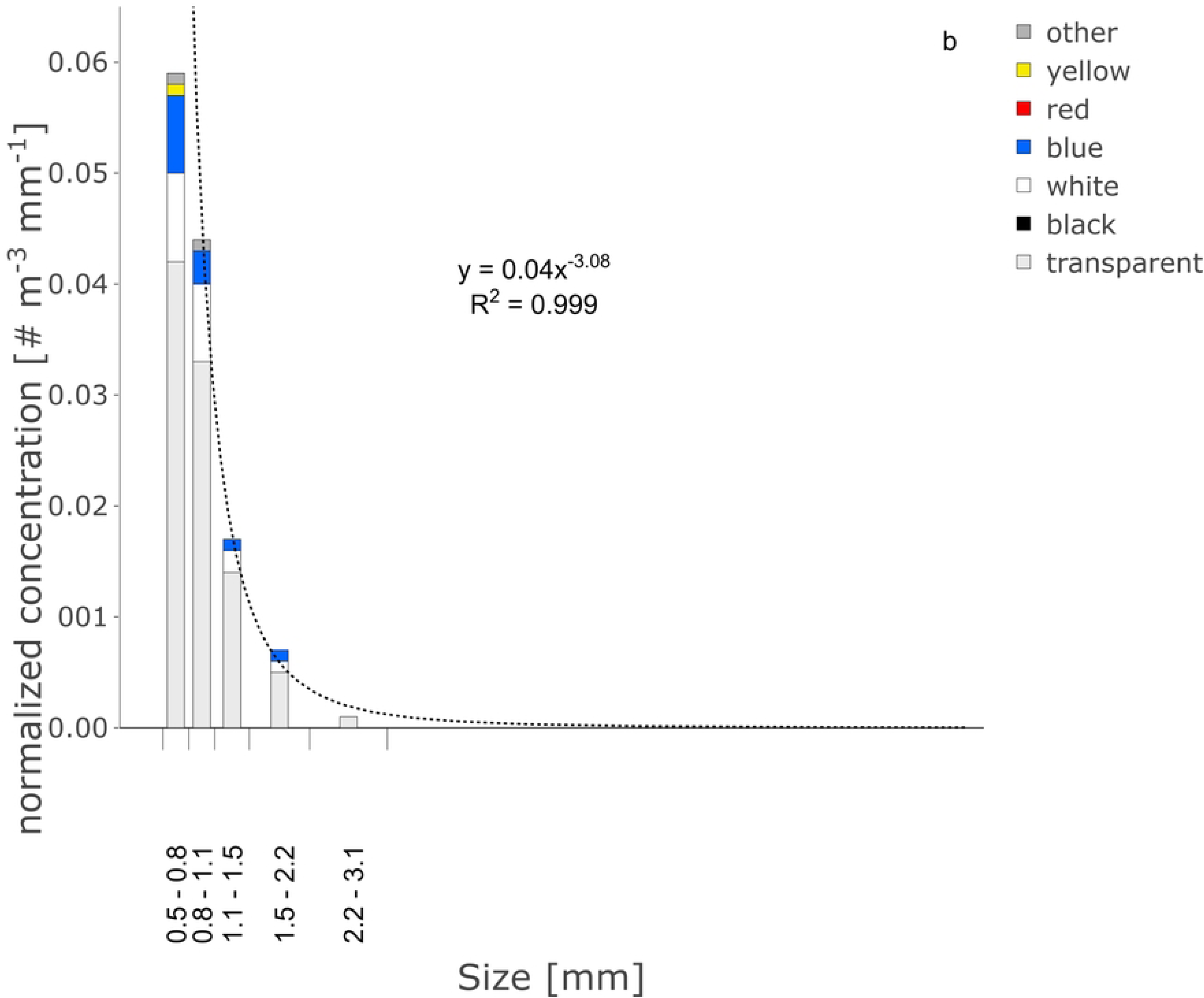
Differential size distribution of (a) fibers and (b) fragments depicting contribution of colors in each size bin. The best power-law fit is also shown. The smallest size bin was ignored for the calculation of the power law fit, as it is likely under-sampled.

The differential size distribution of probable microplastic particles revealed a strong power law fit for both fibers (*A* = 0.109, *b* = 1.87; r^2^ = 0.997, n = 4, p < 0.01) and fragments (*A* = 0.039, *b* = 3.08; r^2^ = 0.999, n=4, p < 0.01) (**Figure 4**). The low *b* value for fibers indicates a high proportion of long fibers within the measured size range.

### 3.2 Carbon content of marine particles and microplastic

Particulate Organic Carbon > 300 μm ranged from ≤ 0.06 mg C m^-3^ (stations S5, S8, S9) to > 2.6 mg C m^-3^ (stations S6, S7) with a C: N ratio of 5.4 to 6.2 (mol: mol) (**Table S2**). The comparison between the measured carbon mass of organic material, including microplastic, with that of the calculated probable microplastic carbon mass, revealed that microplastic contributed to POC < 5% at stations S5, S6, and S7, and ≥ 13% at stations S8 and S9 (**Table S2**).

## 4 Discussion

### 4.1 Microplastic in Placentia Bay

Microplastic concentrations tend to be high in bays and semi-enclosed water bodies near densely populated regions, such as in Tokyo Bay, the Gulf of Mexico, and the Baltic Sea (27, 41, 42), because most microplastic originates from terrestrial anthropogenic sources (43, 44). The average probable concentration of > 300 μm microplastic particles in Placentia Bay was 0.31 ± 0.19 particles m^-3^, which should be considered a conservative estimate, due to our cautious assumptions. This average *probable concentration* of > 300 μm microplastic particles in Placentia Bay is higher than the median of 0.26 particles m^-3^ of 377 surface samples collected in marine waters globally using comparable nets with a mesh size between 300 and 333 μm (**Table S3**). Microplastic concentrations in these 377 samples, which stem from 37 studies (40), ranged over three orders of magnitude from < 0.02 part. m^-3^ (Arctic and Antarctic) to > 100 particles m^-3^ (Sri Lanka, Sea of Korea) (**Table S3**). Concentrations in Placentia Bay were most similar to sites in the Baltic, Adriatic, Bohai, Mediterranean, and Andaman Seas, all areas with relatively high human population densities. It is also higher than even the maximum concentration found in the St Lawrence River system, a much more populated area in Atlantic Canada (45). Thus, our hypothesis that microplastic concentrations in Placentia Bay would be low compared to global averages was not supported.

A comparison to specific bays of similar size, where sample collection (similar nets) and processing (digestion and analysis) were similar, revealed that maximum concentrations in Placentia Bay were roughly equivalent to those found in Chesapeake Bay and the Bay of Biscay, despite their larger sizes and higher human population densities (**Table** 1). Tokyo Bay, which is similar in size to Placentia Bay but has a much higher coastal population density, shows peak microplastic concentrations that are 1–2 orders of magnitude higher (27). The minimum microplastic concentration measured in Placentia Bay is 1-2 orders of magnitude higher than those in Chesapeake Bay and the Bay of Biscay and only eight times lower than those in Tokyo Bay. Thus, while peak concentrations of microplastic in Placentia Bay appear similar, minimum concentrations are high compared to measurements in bays with higher population densities, challenging the expectation of relatively low microplastic concentrations in Placentia Bay. While large differences in microplastic concentrations may also be caused by methodological differences, these comparisons focus on studies using similar methodological approaches, trying to avoid biases.

**Table 1.**
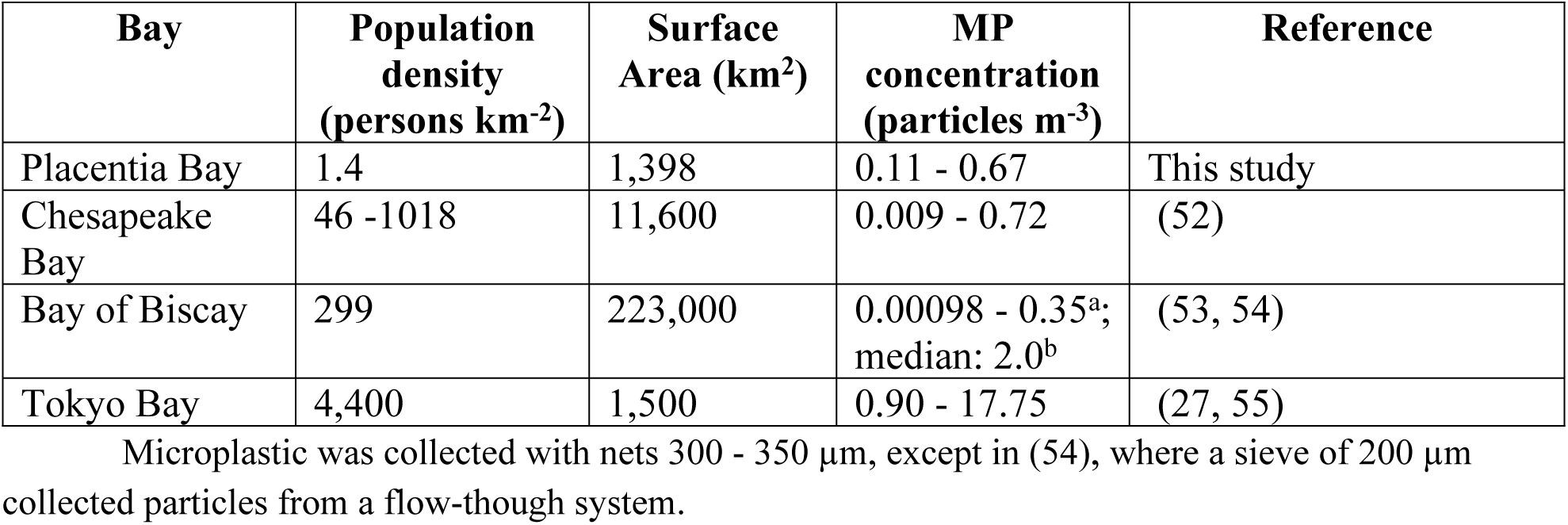
Comparison of > 300 μm microplastic particles in four bays varying in size and coastal human populations.

The contribution of microplastic carbon to the organic carbon captured in 300 µm nets was at the high end (2-14%) of ranges presented in other studies (46, 47): Carbon from smaller microplastics particles (1-300 μm) are estimated to contribute 1–3% of measured POC to New York Bight surface waters (48), between 0.3% and 2.1% to surface waters of the Pacific (Zhao et al. 2023) and 5% at depth, where non-plastic POC concentrations are low (49). Microplastic contribution to vertical POC flux as measured with sediment traps is estimated to contribute on average 1.5% to POC (50). The high values measured in Placentia Bay only consider particles > 300 μm in size, excluding most phytoplankton, likely explaining the larger contribution of microplastic to POC. This suggests that the negative impact of microplastic on filter- and suspension-feeders may be felt more by predators selecting for larger particles, due to the decreased availability of digestible and degradable organic carbon (47). Significant contribution of microplastic to organic carbon will also have, notable impacts on ecosystem budgets and predictions of carbon cycling (51), as well as interfere with ^14^C-based age determinations (49).

### 4.2 Spatial Distribution

The spatial distribution of microplastics within Placentia Bay can largely be explained by water circulation, as there is no significant river discharge into the Bay (28). A three-dimensional circulation model driven and validated by observational data, reveals that while the mean surface circulation generally follows a cyclonic (counter clockwise) pattern, southwesterly winds induce coastal upwelling in summer and early fall (28), when sampling took place. During this time, surface circulation within the Bay is eastward (28). In accordance with these current patterns that would push floating particles against the eastern shore (28), higher microplastic concentrations were observed in the eastern Bay (e.g., S8) compared to the western Bay. The inner Bay’s longer water residence times compared to the outer Bay (28), which is open to the Atlantic, may explain the relatively high microplastic concentrations at inner Bay stations compared to those in the outer Bay. (28). The observed microplastic distribution patterns also align with the distribution of cod eggs, which at this time of the year tend to concentrate in the inner and eastern parts of the Bay (56), suggesting that currents and residence time control distributions of both particle types.

Potential terrestrial point sources (**Fig. 1**) do not directly correspond to microplastic concentrations at inshore sampling location (**Fig. 2**); except for a possible link between the elevated concentrations at station S1 and a cluster of potential source sites near Placentia. The large difference between replicate net tows (**Fig. B1**) at this station indicates that possibly an isolated patch was sampled with one net. Similarly, point sources could not explain microplastic abundance in the St Lawrence River and Estuary, although source sites in rivers are often closer to points of discharge and flow is mostly one directional (45). In Placentia Bay, conceivably currents masked potential signals from terrestrial point sources with one exception: Arnold’s Cove beach (47°46.5’N, 53°58.3’W) located in the inner Bay, east of station S7 and north of station S8, was identified as a hot spot of plastic pollution during a shoreline plastic debris survey (57). Beaches frequently act as temporary reservoirs and effective microplastic source sites (10). Plastic shoreline litter at Arnold’s Cove consisted of 60% fishing gear (57), and likely contributed to the high microplastic concentrations at station 7 and especially station 8, which in summer is downstream from Arnold’s Cove.

### 4.3 Tracking microplastic sources

The low diversity of microplastics in Placentia Bay, where nearly half of all microplastics were long, transparent, PP fibers, is conspicuous (16, 36, 44, 58–60). Globally, most (75%) synthetic fibers in ocean microplastic surveys are thought to consist of PE, with nylon fibers contributing 11%, and PP of any color and size only 6% (61). However, PP particles are more important in nearshore waters, whereas offshore PE overshadows PP, possibly due to a faster abiotic loss of PP (49). Nevertheless, a comprehensive study of microplastic in net tows (> 300 μm) in the Mediterranean Sea found PP to contribute between about 5% and 25% to microplastic inventories (20), and a metaanalysis of 24 Bays also found that PP contributed on average 25% to all microplastic particles (19). Thus, a contribution of PP of over 60% may be considered unusual and noteworthy.

The dominant shape and color of microplastic may also vary widely, with, for example, fragments dominating (75%) in Tokyo Bay (27), but fibers contributing > 90% in the Bay of Biscay (54). And while clear, white and green colors were most important in Tokyo Bay (27), blue, gray and orange dominated in Chesapeake Bay (52). This variability notwithstanding, it is highly unusual to see one type of particle as defined by all three characteristics, color, polymer, and shape, contribute nearly 50% of all particles in microplastic inventories, making our observation of the dominance of transparent PP fibers in Placentia Bay exceptional.

Additionally, the low *b*-value of the size frequency distribution of fibers in the Bay implies that long fibers contributed disproportionately, consistent with a recent influx of long PP fibers, rather than a continuous, long term, low-level release. The faster fragmentation rate of PP fibers compared to PE fibers commonly leads to the opposite effect, where small PP fibers dominate the size distributions of PP fibers (20).

The unique, specific composition of microplastic particles in Placentia Bay, an area where continuous input from common sources of fibers, such as tires or textiles, are low, suggests that a distinctive input event was responsible for these long PP fibers. PP fibers are utilized in a variety of materials, including thermal insulation, sheets, blankets, clothing, performance gear, carpets, and bags (62). Transparent PP fibers are also used in the production of fiber reinforced concrete, to improve the strength and flexibility of the concrete (63, 64). And transparent PP fibers are the main weathering product of medical face masks (21).

A concrete manufacturing plant is situated along the coastline of Placentia Bay (**Figure 1**), and an accidental spill of such fibers could be a potential explanation. The PP fibers added to concrete as reinforcement are frequently 12 mm long, but would rapidly fragment to the size of microplastic when released to the ocean. We found no report of an accidental release of such fibers.

Another potential source are disposable facemasks (e.g., N95, medical surgical masks), widely used during the COVID-19 pandemic. When they deteriorate, the vast majority of these masks, independent of brand or design, shed mostly (> 80%) relatively long (> 100 to 300 µm), transparent PP fibers (23–26). When masks disintegrate, they also form fiber aggregates or balls, like the ones we observed. A back-of-the-envelope calculation confirms that transparent PP fibers in Placentia Bay could realistically stem from disposable facemasks released into the Bay in the 1 - 1.5 years before sample collection. This order of magnitude estimate (**Table** 2) is based on (#1) the amount of transparent PP fibers in Placentia Bay, which was calculated from our measured, average concentration and the area of the Bay; (#2) the number of masks that entered the Bay estimated from population density, assumed mask usage and the fraction of masks discarded into the Bay; and (#3) release rate of transparent PP fibers from medical facemasks as measured experimentally. Using an exponentially decreasing release rate of fibers, and assuming an average of 180 days of weathering per mask, the number of masks needed to explain the observed amounts of transparent PP fibers in Placentia Bay (#4) may be calculated and compared to the estimate of masks entering the Bay (#2; **Table** 2). The largest uncertainty in this back-of-the-envelope calculation resides in the approximations of the release rate of fibers from masks.

**Table 2:** Variables considered ascertaining if transparent > 300 μm PP fibers (tr-PP-F) in Placentia Bay (PB) may originate from degrading medical facemasks released to the ocean in the previous year (12 months’ timespan).

| # | Estimated Values | Facts & Numbers | Assumptions | Justifications |
| --- | --- | --- | --- | --- |
| **1** | Number of tr-PP-F in PB in summer 2021: $\sim 10^9$ tr-PP-F | Probable concentration of tr-PP-F<br>Area of PB (1398 m<sup>2</sup>): | Average concentration is homogenous in the upper 5 m, and zero below 5 m | This input event would be recent and a distribution of tr-PP-F to all depths would not be expected. |
| **2** | Number of facemasks that entered the ocean based on population and usage: 19,000 to 62,000 masks | Coastal population (Marystown, Placentia, and Burin): 12,000 people. This is likely an underestimate | Average weekly usage summer 2020-21: 1 mask per person, total $0.6 * 10^6$ masks.<br>3% - 10% of discarded masks enter the ocean (68): 3% common, 10% high estimate | Before summer 2020 medical facemasks were rarely available. Thereafter, children often did not use masks, but health facilities used a lot.<br>Washing medical masks creates > 48,000 fibers > 0.5 mm) (26).<br>1 mask per week per person after summer 2020 represents an unbiased average. |
| **3** | Release rates of tr-PP-F > 300 µm:<br>Including UV exposure: $\sim 10,000 \text{ d}^{-1}$ (68)<br>Including mechanical abrasion (beaches): $> 35,000 \text{ d}^{-1}$ (24)<br>At those rates, > 35,000 and > 50,000 fibers > 300 µm are released per mask within 180 days. | Rates depending on weathering (25), (23, 68, 70) (24)<br>Time series data was extrapolated to 180 days, assuming an exponential decrease in the release rate | Release rate decreases exponentially over time (23).<br>Masks exposed to UV (68) or mechanical abrasion on shoreline (24) | The assumed release rates excluded the extremely high and low release rates observed in laboratory experiments and are lower than the averages calculated from (23, 24, 65, 68). They are considered conservative |
| **4** | Weathering of masks that would create $\sim 10^9$ tr-PP-F > 300 µm: 20,000 to 28,500 masks, depending on release rate (see #3) | NA | NA | The estimated number of masks need to explain the observed tr-PP-F (#4) are at the low end of our estimates of masks that likely entered the Bay (19,000 to 62,000); (see #2). |

The total number of fibers released per mask during its lifetime is thought to be > 0.9-1.5 * 10^6^ or even > 10^8^ (26, 65, 66), but full deterioration to microplastic is thought to take 15 years (67).

However, laboratory experiments reveal that artificial weathering with UV light turns PP brittle and accelerates the initial fragmentation rate drastically, compared to mixing alone (68, 69). Initial release rates of > 300 µm PP fibers, after UV exposure (∼10,000 fibers d^-1^) (68) and assuming an exponential decrease in release rate (23) suggest that about 28,000 masks would explain the transparent PP fibers in Placentia Bay. This estimate lies within the lower range of masks predicted to have entered the ocean (#2, **Table 2**) and thus could explain 100% of the transparent PP fibers present in the Bay. If weathering due to mechanical abrasion with sand is taken into account, the initial release rates of transparent PP fibers would be higher, > 35,000 fibers d^-1^ (24), and 20,000 masks entering Placentia Bay could easily explain our transparent PP fiber concentrations, even assuming that only 3% of discarded masks (not 10%) entered the Bay (**Table 2)**. Experimentally aging *in situ* for two months led to the rapid disintegration of masks and the release of 6 * 10^8^ fibers of which > 20 % were > 0.5 mm long (26), suggesting even higher release rates. These varied estimates demonstrate that it is possible that many of the transparent PP fibers detected in Placentia Bay originated from discarded medical facemasks used during the pandemic. Similar input events would be expected in other aquatic systems worldwide, but may not be easily detected if continuous input rates of diverse microplastic are high.

Assuming an input event of transparent PP fibers skewed the composition of microplastic in the Bay, it is worthwhile to focus on the remaining microplastic particles in our inventories.

Aside from the transparent PP fibers, microplastic composition in the Bay were much more diverse, with transparent and white PE fragments and blue PP fibers contributing most. These microplastics are likely to be predominantly attributed to fishing activity, with small contributions from land-based sources providing the remainder. Tangled fiber balls observed at multiple sites also resembled those generated from beached fishing gear (15) and during rope abrasion experiments (13). Fishing pots used in Newfoundland, commonly use green or blue PE netting (71), a possible source of some of the PE fibers found in our study. Nets are commonly white or transparent PE, PP or nylon. It is noteworthy that no nylon fibers were found during our study, although nylon gill nets were introduced to the Newfoundland cod fishery in the early 1960’s (72) and commonly used because they were subsidized by the Canadian government (73). It is estimated that > 80,000 gillnets were lost in Atlantic Canadian waters between 1982 and 1992. However, nylon is denser than seawater, and nylon fibers may have largely accumulated in sediments.

## Summary and Conclusions

While the precise origin of the microplastic observed in Placentia Bay remains uncertain, their specific characteristics are consistent with the hypothesis that the transparent PP fibers observed in 2021 largely originated from disposable facemasks, providing nearly half of the microplastic in the Bay. Since medical face masks were used widely during 2020/ 2021, a distinct signal of PP fibers may be expected to appear in diverse aquatic systems. However, such a pulsed input of transparent PP fibers stemming from medical facemasks, would likely get lost in bays with a large, continuous, terrestrial-sourced input of microplastic, as the “signal to noise ratio” would be low. The generally low microplastic inputs into Placentia Bay due to low anthropogenic activity, and the lack of appreciable river discharge may have made such an input pulse visible. Transparent and white PE fragments and blue PP fibers likely arose from fishing activity predominantly, with additional input from land-based sources associated with coastal communities resulting in the higher diversity of the other half of microplastic inventories observed in the Bay.

The distribution of microplastic within the Bay was primarily driven by currents and residence time of water masses, rather than the location of land-based sources, with one exception. Microplastic concentrations were elevated downstream of a beach, which acted as a temporary reservoir for plastic detritus. This finding suggests that clean-up of plastic on beaches has positive effects on the surrounding waters. The absence of links between potential source sites and waterborne microplastic implies that the rate of microplastic formation from macroplastic detritus is frequently slow compared to the timescales of microplastics particle transport by water movements, e.g., microplastic distribution in bays is often dominated by hydrography, not locations of source sites. Consequently, potential source sites cannot necessarily be identified based on their proximity to the microplastic. This study is relevant beyond Placentia Bay, as similar microplastic input events likely occurred globally during the pandemic. The Bay’s unique characteristics make it an ideal natural laboratory for observing and analyzing such an event.

## Acknowledgements

We thank Vonda Hayes for discussion on fishery practices in the sixties. All Raman spectral data were produced in SBU’s School of Marine and Atmospheric Sciences’ NAno Raman Molecular Imaging Laboratory (NARMIL), a community resource dedicated to environmental science applications and founded with NSF-MRI grant OCE-1336724. We thank for funding the Ecosystems and Oceans Science Contribution Framework, EOSCF, Department Oceans and Fisheries, as well as the Canada Research Chair Program to UP. The support of the Natural Sciences and Engineering Research Council of Canada (NSERC) for the Discovery Grant and the Collaborative Research and Training Experience Program, PEOPLE (Persistent, Emerging, and Oil PoLlution in cold marine Environments) is also appreciated. All data is available from the St. Lawrence Global Observatory data catalog https://doi.org/10.26071/10.26071/02c0aa6b-2043-4461

## Supporting Information captions

<u>Table S1</u>: Location of sampling stations

<u>Table S2</u>: Contribution of Microplastic to Particulate Organic Carbon

<u>Table S3</u>: Concentrations of Microplastic in Global Surface Waters Collected with 300 µm to 350 µm Nets

<u>Figure S1</u>: Comparison of Net Types: Uncorrected microplastic concentrations (particles m^-3^) for fibers and fragments at each station as collected with the plankton (Pl.) or neuston (Ne.) nets and determined by visual identification

<u>Figure S2</u>: Calculating Microplastic Concentrations: Diagram depicting corrections implemented to estimate microplastic concentrations from visual counts corrected with insights gained via Raman microspectroscopy

## Notes

### Competing Interest Statement

The authors have declared no competing interest.

